# Machine Learning Strategies for Improved Phenotype Prediction in Underrepresented Populations

**DOI:** 10.1101/2023.10.12.561949

**Authors:** David Bonet, May Levin, Daniel Mas Montserrat, Alexander G. Ioannidis

## Abstract

Precision medicine models often perform better for populations of European ancestry due to the over-representation of this group in the genomic datasets and large-scale biobanks from which the models are constructed. As a result, prediction models may misrepresent or provide less accurate treatment recommendations for underrepresented populations, contributing to health disparities. This study introduces an adaptable machine learning toolkit that integrates multiple existing methodologies and novel techniques to enhance the prediction accuracy for underrepresented populations in genomic datasets. By leveraging machine learning techniques, including gradient boosting and automated methods, coupled with novel population-conditional re-sampling techniques, our method significantly improves the phenotypic prediction from single nucleotide polymorphism (SNP) data for diverse populations. We evaluate our approach using the UK Biobank, which is composed primarily of British individuals with European ancestry, and a minority representation of groups with Asian and African ancestry. Performance metrics demonstrate substantial improvements in phenotype prediction for underrepresented groups, achieving prediction accuracy comparable to that of the majority group. This approach represents a significant step towards improving prediction accuracy amidst current dataset diversity challenges. By integrating a tailored pipeline, our approach fosters more equitable validity and utility of statistical genetics methods, paving the way for more inclusive models and outcomes.

## 1. Introduction

In recent years, genome-wide association studies (GWAS) have provided many insights into the genetic basis of complex traits and diseases. However, these findings predominantly benefit populations of European descent due to their over-representation in genomic datasets. Individuals with Asian, African, and other ancestries only represent a small fraction of the available datasets.^1^ Although individuals of European descent constitute *∼*79% of GWAS participants,^2^ they account for less than a quarter of the global population. This disproportionate representation creates a limitation in precision medicine, because statistical models built to infer disease risks or health-related traits can perform poorly for individuals from populations that were underrepresented when creating the model, exacerbating health disparities. Despite initiatives to include a broader range of populations in genetic studies and biobanks,^3–7^ the proportion of non-European individuals in GWAS studies has stagnated in the last decade.^2,8^ This imbalance has a direct impact on Polygenic Risk Score (PRS) prediction for underrepresented populations,^9^ making clinical applications based on PRS significantly more accurate for individuals of European descent, but less effective for other populations.^10–12^ This disparity has raised ethical concerns within the scientific and clinical community.^1,3,8,13^ While most studies only use European individuals and European-derived statistics to build predictive models,^8,11,14^ recent studies have explored including non-European training data in PRS construction, but this has only proven effective when a large number of training samples of non-European target populations are available.^15^

Phenotype prediction utilizes genetic information to forecast an organism’s observable characteristics, known as phenotypes. These traits can range from disease susceptibility to other attributes, enabling personalized treatments based on individual genetic profiles. Machine learning (ML) and deep learning (DL) models used to predict phenotype and population structure from genomic data^14,16–20^ are similarly negatively impacted by imbalanced datasets. Vokinger et al.^21^ highlighted the presence of bias in ML-based medicine prediction pipelines. Specifically, they revealed how a naive application of simple ML methods can showcase an overall good performance, yet still produce biased predictions favoring the majority population at the cost of lower accuracy for underrepresented groups. Efforts to mitigate this bias exist, such as Afrose et al.^22^ who created a double prioritized bias correction technique that involves training customized prediction models for specific subpopulations. However, this approach is limited to binary classification tasks and is not generalizable to other prediction problems.

Conventionally, the statistical methods that are applied for genomic prediction problems linearly combine the effects of different genetic variants on an individual’s risk of disease. Some of the most widely used regression models include Lasso,^23^ a linear method with *𝓁*_1_ penalty, Elastic net^24^ with *𝓁*_1_ and *𝓁*_2_ penalty, and efficient implementations of both.^14^ Although being the routine choice in most studies, linear models are not able to capture non-linear genetic interactions that can contribute to a phenotype.^25^ The ability of non-linear predictive models to capture genetic interactions could help improve performance generalization across populations.^26,27^ Neural networks, a complex non-linear method, have recently gained traction in computational biology,^28,29^ but require vast amounts of data for training. Large-scale biobanks, such as the UK Biobank,^30^ provide such expansive datasets. However, the small proportion of samples from minority populations hinders robust generalization across different genetic backgrounds. In contrast, gradient boosting (GB) algorithms,^31^ such as eXtreme Gradient Boosting (XGBoost)^32^ and LightGBM,^33^ have frequently demonstrated superior performance for tabular data and small-sized datasets,^34,35^ and have already been explored in biological studies for tasks such as local ancestry inference,^36^ protein-protein interactions,^37^ and drug-gene interactions.^38^ In the realm of genotype-to-phenotype prediction, recent research has also highlighted the potential benefits of using such nonlinear predictive models.^39,40^

In this paper, we aim to improve phenotype prediction for diverse and underrepresented populations. We propose a more inclusive genomic research approach that uses multi-ancestry data together with advanced machine learning techniques to boost the predictability of complex traits across a broader range of populations. Our method leverages several machine learning techniques such as boosting, and ensembling, and we propose population-conditional weighting and re-sampling techniques to generate more accurate models for underrepresented populations without requiring large sample sizes of non-European training data. Fig. 1 illustrates the workflow of our approach, starting with the formation of the data set through the application of various machine learning techniques and data de-biasing methods. We compare our approach with state-of-the-art statistical genetics models on the UK Biobank, conducting a systematic evaluation across 12 phenotypes in European (British), African, East Asian, and South Asian individuals. Given that the majority population is of European descent, we observe a large gap in phenotype prediction accuracy for minority populations when using classical linear methods. This disparity only grows when European-only data is used to train any of the prediction models. We demonstrate how the application of our method helps narrow this accuracy gap, balance the performance across populations, and obtain state-of-the-art phenotype prediction results for multi-ancestry datasets.

**Fig. 1.**
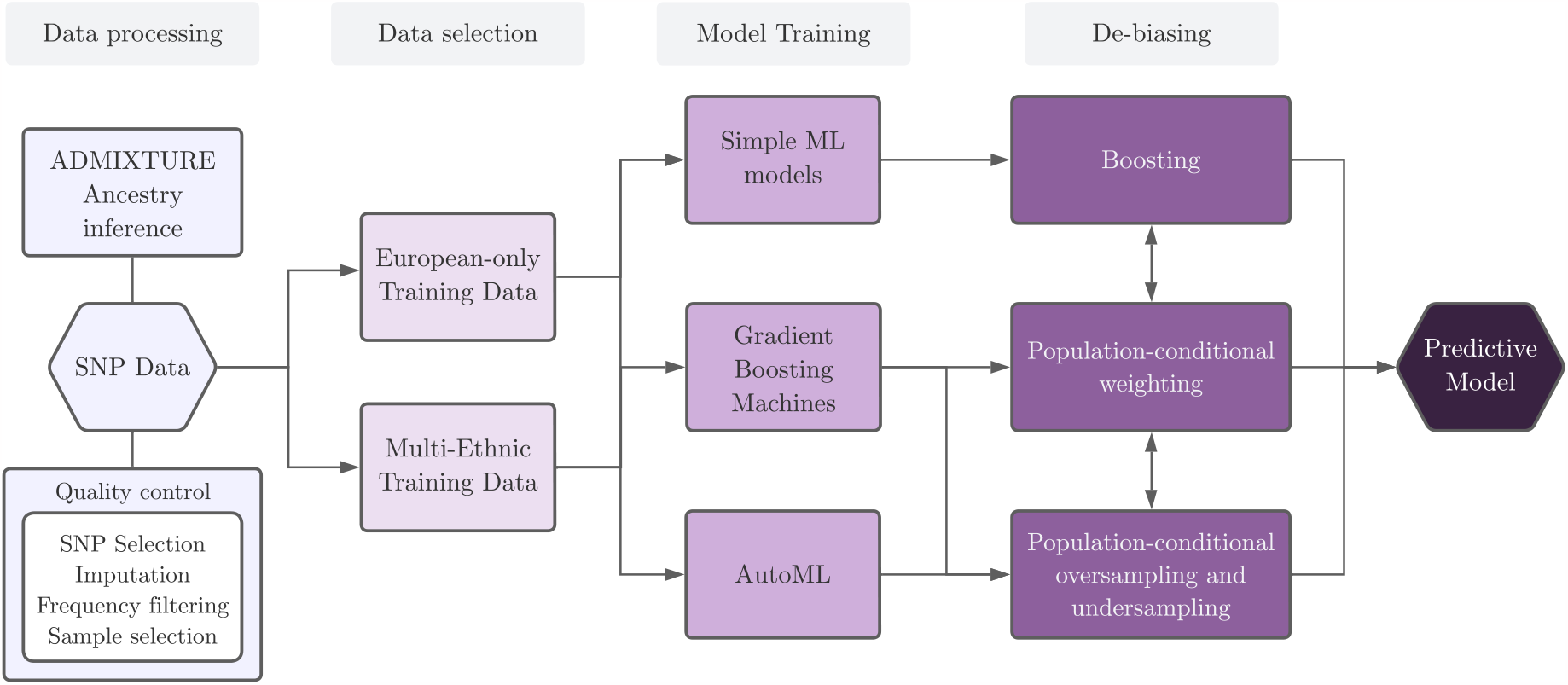
A schematic representation of our predictive modeling pipeline, starting from the initial data ingestion to the application of various ML methods and de-biasing techniques.

## 2. Methods

### 2.1. Dataset preparation

We utilize a dataset extracted from the UK Biobank^30^ that includes European (British), South Asian, African, and East Asian individuals (see Fig. 2). We use the pre-computed population labels from the Global Biobank Engine (GBE),^41^ inferred based on genetic clustering with ADMIXTURE software^42^ results, which provides a maximum likelihood estimation of an individual’s genetic ancestry clustering from multilocus genotype datasets.

**Fig. 2.**
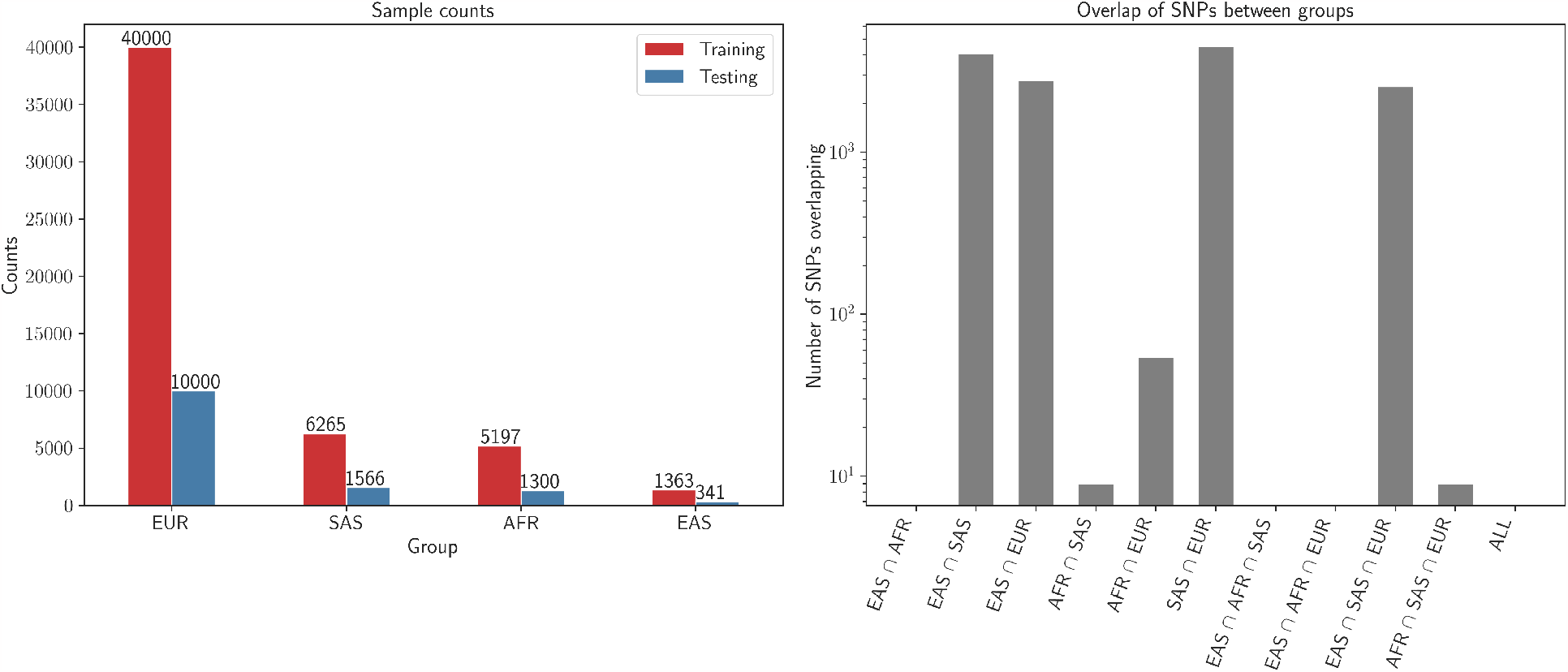
EUR - European British, SAS - South Asian, AFR - African, EAS - East Asian (Left) Sample counts per group in the training and testing set. (Right) Percentage of SNP overlap between the selected sets of SNPs per group using the MAF filter.

Single nucleotide polymorphism (SNP) sequences are encoded using a ternary system, where at each genomic position, an individual *i* has *n*_*ij*_ *∈* {0, 1, 2} copies of the minority SNP *j*. To address high dimensionality and retain the most informative SNPs, we apply a SNP selection process. Minor allele frequency (MAF) filtering is applied with a 1.25% threshold, keeping a set *𝒮*_*p*_ of 10000 SNPs for each population *p ∈ P*, such that | *𝒮*_*p*_| = 10000. After SNP selection for each population, we computed the union of these sets. It is important to note that not all sets necessarily overlap with every other set. The union is represented by:

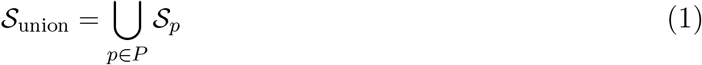

This resulted in a unified set of SNPs where |*𝒮*_union_| = 31153, which is then used for all individuals, creating a dataset of 66032 individuals and 31153 features. Fig. 2 shows the intersection size of the sets of selected SNPs *𝒮*_*p*_ for all intersections of populations *p ∈ P*. We observe that the highest overlap is between South Asian, East Asian, and European populations, while the selected set of SNPs for the African population has practically no overlap with the others. Any subsequent missing SNPs within the samples underwent mode imputation to ensure data completeness.

To conduct our experiments, we study a set of phenotypes included in the GBE,^41^ listed in Table 1. Details regarding the correspondence of the GBE to the UK Biobank can be found in the GBE paper. We selected the available phenotypes with minimal missing data for the minority populations, and that also showed good predictive performance from genotype features.^43^ We analyze both binary phenotypes (absence or presence of the phenotype) and continuous phenotypes to evaluate model performance across both classification and regression tasks.

**Table 1.**
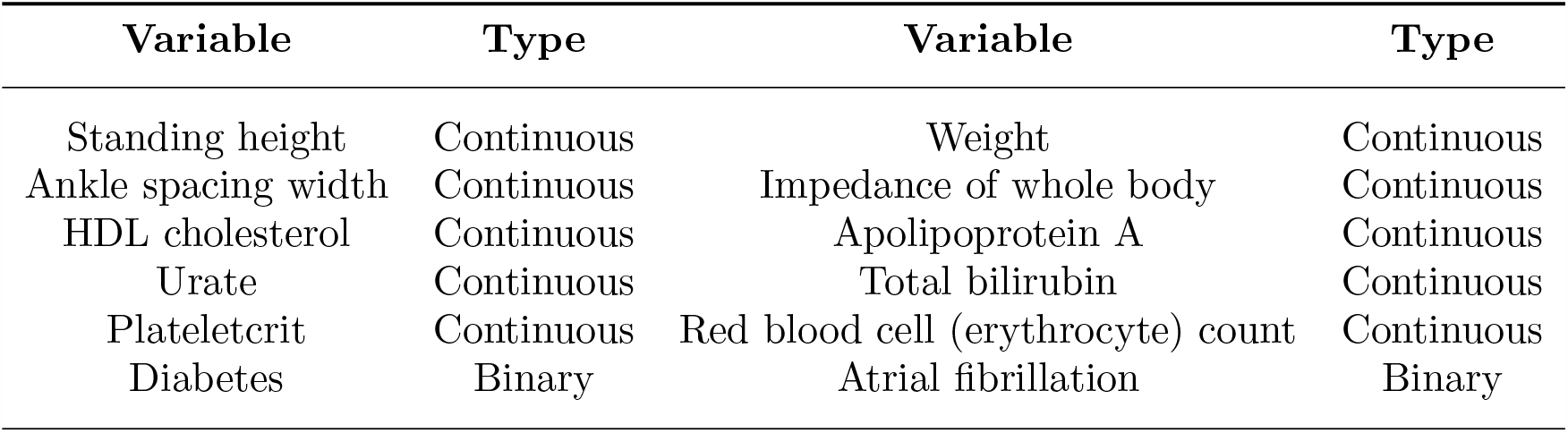
We present results on 12 phenotypes, 10 continuous and 2 binary ones.

| Variable | Type | Variable | Type |
| --- | --- | --- | --- |
| Standing height | Continuous | Weight | Continuous |
| Ankle spacing width | Continuous | Impedance of whole body | Continuous |
| HDL cholesterol | Continuous | Apolipoprotein A | Continuous |
| Urate | Continuous | Total bilirubin | Continuous |
| Plateletcrit | Continuous | Red blood cell (erythrocyte) count | Continuous |
| Diabetes | Binary | Atrial fibrillation | Binary |

Additionally, we ensured there is no missing data and filter samples that have missing phenotypic information. The dataset is partitioned into a training set and a testing set, comprising 80% and 20% of the data, respectively. We applied stratified sampling, ensuring the proportion of samples from each population closely mirrors their proportion in the overall dataset.

### 2.2. Algorithmic models

We explore a wide range of machine learning methods to improve phenotype prediction on underrepresented populations. Some algorithms serve as standalone models, capable of making predictions without supplementary techniques. Other algorithms we describe in this section, such as boosting, are techniques that can be used to further improve the performance of a base machine learning model. Finally, we explore complex machine learning systems that combine multiple models and automate the process of machine learning.

#### Linear models

We include the Least Absolute Shrinkage and Selection Operator (Lasso),^23^ a linear regression method that performs variable selection (i.e., identifies the most important predictors) and regularization, which prevents overfitting by constraining the model parameters. It does this by imposing an *𝓁*_1_ penalty, effectively reducing some coefficients to zero. We also use Elastic Net,^24^ a regularized method that combines *𝓁*_1_ and *𝓁*_2_ penalties, allowing coefficient shrinkage and feature selection.

#### Boosting

We consider boosting,^44^ a powerful ensemble machine learning technique that constructs a strong predictive model by combining multiple *weak learners*—simple models— that are trained sequentially. In each iteration of the boosting process, a new weak learner is trained giving more importance to the instances that were poorly predicted by the previous models, meaning the model attempts to correct the errors of its predecessors. This procedure is repeated sequentially, with each new model targeting the instances where the combined ensemble has performed the worst. The final model is a weighted combination of all the weak models, which often yields a strong predictive performance by aggregating the strengths of all individual models. Decision trees are the most common type of weak learners used in boosting algorithms. However, we also study how boosting can help improve predictive performance when traditional linear methods used in the field, such as Elastic Net, are used as weak learners.

#### Gradient boosting machines

A specific implementation of the boosting techniques are gradient boosting machines (GBM). The key idea behind GBMs is the use of the gradient descent algorithm to minimize a loss function, which quantifies how well the model predicts the target variable. In each iteration, rather than directly focusing on the poorly predicted instances, a new decision tree is fit to the negative gradient (residuals) of the loss function with respect to the prediction of the ensemble model from the previous stage. This new decision tree provides a direction in which the prediction should be adjusted to minimize the loss function. The predictions are then updated by taking a step in this direction. Extreme Gradient Boosting (XGBoost)^32^ and LightGBM^33^ are two optimized implementations of GBMs that have gained significant popularity due to their efficiency and performance. XGBoost offers several advanced features such as regularized boosting, handling of missing values, and tree-pruning that makes it faster and more robust. LightGBM also offers high performance and efficiency but is particularly notable for its effectiveness with large datasets and high-dimensional data, due to its innovative histogram-based algorithm that reduces memory usage and speeds up training.

#### AutoML

Automated Machine Learning (AutoML)^45^ refers to the automated process of end-to-end model development, encompassing steps from feature engineering to model selection, hyperparameter tuning, and model evaluation. AutoML methods have been developed to streamline the machine learning pipeline while reducing time and expertise required to develop effective predictive models. In particular, we consider AutoGluon^46^ (AG), a state-of-the-art AutoML framework known for its robust performance, efficiency and ease of use. AutoGluon automatically trains and optimizes multiple models such as neural networks, nearest neighbors, linear models, and gradient boosting machines, combining them into a stacked ensemble.

### 2.3. Population-conditional re-sampling solutions

We introduce a set of population-conditional re-sampling techniques to address population imbalance in datasets. These techniques serve as auxiliary methods designed to reduce model bias towards the majority population and can be integrated with any predictive model. While we focus on human populations in this work, these techniques can also be applied to any data where samples can be grouped into different populations, groups, or categories. Moreover, they are suitable for tasks beyond single-target classification, such as regression, and they can also be extended to multi-output tasks.

#### Population-conditional oversampling and undersampling

We modify the traditional oversampling and undersampling techniques used in imbalanced classification tasks, and adapt them to address imbalances at the population level, regardless of the target variables (both categorical and continuous). We organize the training dataset as ***X*** *∈* ℝ^*N ×d*^ such that each row represents an individual, and the target variable or variables are concatenated to the rest of the input features as the final attributes. The population label is then used as a downstream label ***y***^*′*^ *∈* ℝ^*N*^ for the oversampling or undersampling rule, originally designed to work with single-target imbalanced classification datasets, such that the “minority” samples are those pertaining to the populations with lowest representation in the dataset. After this procedure, we discard the population labels and split the columns of the re-sampled training dataset as features and targets and fit the prediction models.

We explore population-conditional random oversampling (OS) by picking samples at random with replacement from the minority populations. We also adapt the Synthetic Minority Over-sampling Technique (SMOTE),^47^ which is commonly used to address class imbalances by generating synthetic samples. Our modification enables us to synthetically increase the number of instances from the minority populations in the training set. Note that in the case of regression tasks, our approach differs from existing SMOTE variations for regression,^48,49^ which involve identifying “minority” samples based on the distribution of the target values rather than external categorical labels associated with the samples. Finally, we also consider adapting the SMOTE-Edited Nearest Neighbours (SMOTE-ENN) algorithm,^50^ a method that combines both oversampling and undersampling techniques. Our proposed population-conditional variation can also be applied to any other re-sampling technique originally designed to address class imbalance in classification problems.

#### Population-conditional weighting

In a similar fashion, traditional class-based sample weighting techniques for class imbalance give more importance to underrepresented classes in the target variable. In contrast, we propose to emphasize the individual instances from underrepresented populations given the population labels each sample has assigned, regardless of their target variable. We calculate *N*_*p*_, the size (i.e. number of samples) of each population *p ∈ P* in the training set, and assign a weight 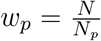 to each sample corresponding to population *p*, inversely proportional to the size of its population, where *N* is the total size of the training dataset.

### 2.4. Evaluation setup

For training, data is either filtered to only contain European ancestry individuals, mirroring the typical bias seen in many genetic studies, or contain the complete, multi-ethnic dataset that includes individuals from underrepresented populations. The testing data is fixed and contains samples from each population group, allowing the assessment and model performance comparison across each population in all the experiments. Model hyperparameters are adjusted by 5-fold cross validation, with hyperparameter configurations drawn from comprehensive search spaces until 1000 configurations are explored or a search budget of 120 hours is reached. Then, the model is fitted on the full training set with the chosen hyperparameter configuration, and evaluated on the held out test set (20% of the data).

Predictive performance is evaluated using the coefficient of determination (*R*^2^) for regression tasks, and the Area Under the Receiver Operating Characteristic Curve (ROC AUC) for classification tasks. *R*^2^ represents the proportion of variance in the predicted phenotype that is explained by the genotype, and its value lies between 0 and 1. An *R*^2^ nearing 1 signifies the model’s high accuracy in phenotype prediction using the given genetic data. In contrast, values approaching 0 highlight the model’s limited predictive capability. ROC AUC measures the model’s ability to distinguish between the positive and negative classes. The value ranges from 0 to 1, with 0.5 indicating performance equivalent to random chance, and values approaching 1 indicating high predictive accuracy.

## 3. Results

### 3.1. Continuous phenotypes

We first analyze the use of multi-ethnic data and the predictive performance of several linear and non-linear models, including Lasso, Elastic Net, LightGBM, and XGBoost, for the 10 continuous phenotypes described in Table 1. Fig. 3 shows the increase in *R*^2^ when training the models with multi-ethnic data, compared to training with only with European individuals on a linear model (Lasso), which is the common practice in the field. Note that relative performance (ratio) cannot be computed per population, as the baseline model obtains an *R*^2^ of 0 for some population groups when predicting some of the phenotypes. We observe that prediction performance significantly improves across all populations and methods when including multi-ethnic data in training. Specifically, the gradient boosting method LightGBM is the model that obtains the highest boost in predictive performance consistently across all ancestry backgrounds, including European and underrepresented ones.

**Fig. 3.**
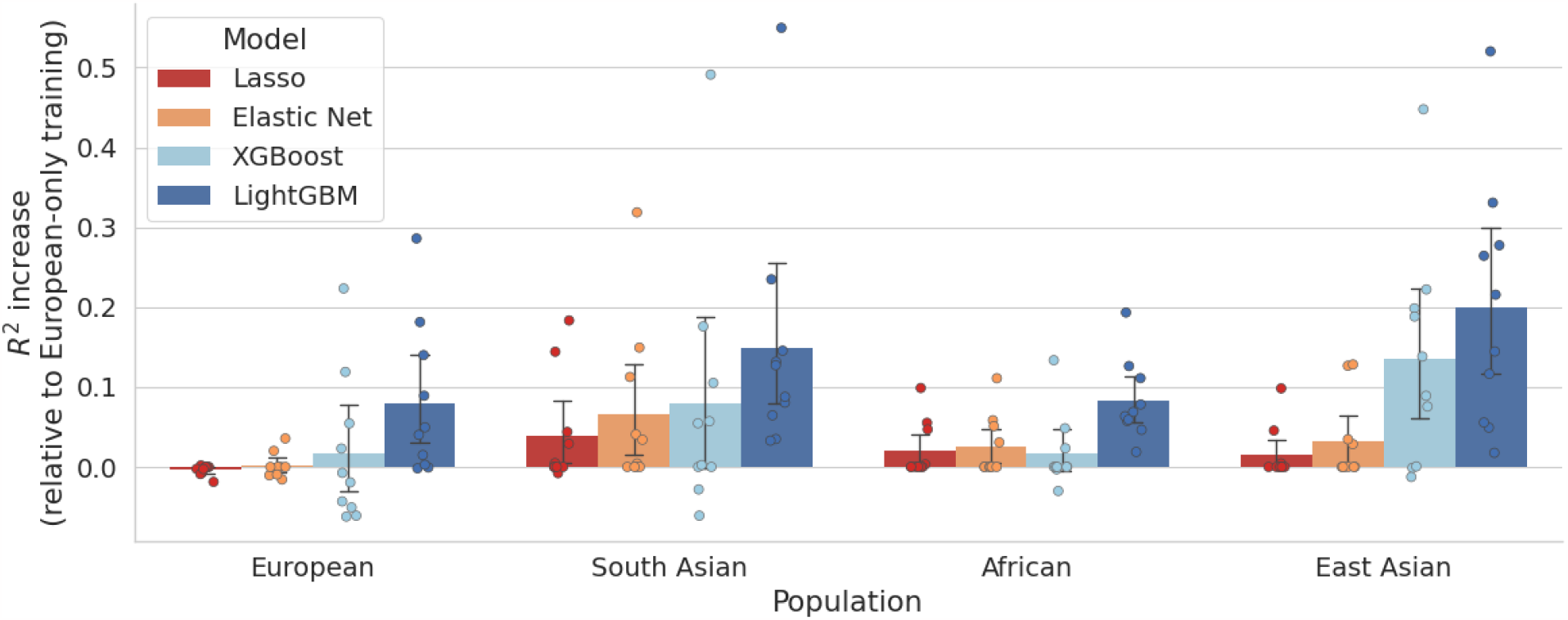
Aggregated results of increase in *R*^2^ for the 10 continuous phenotypes, with a 95% confidence interval, comparing the scores for models trained on multi-ethnic data (including populations underrepresented in the UK Biobank) versus models trained exclusively on the British-with-European-ancestry population.

In an effort to gain deeper insights into how various methodologies can influence a phenotype, we focus on the Standing Height phenotype. Fig. 4 shows our experiments on different models and techniques, with the complexity of machine learning techniques increasing from left to right. Our experiments begin with Elastic Net (EN), starting from a simple linear model trained on individuals of European descent. We then include multi-ethnic data and introduce population-conditional weighting during training. Subsequently, we explore creating an ensemble of Elastic Nets using boosting. As a more complex boosting algorithm, we include LightGBM, followed by AutoGluon, an AutoML method that trains multiple ML models to form a stacked ensemble, including LightGBM as one of its members.

**Fig. 4.**
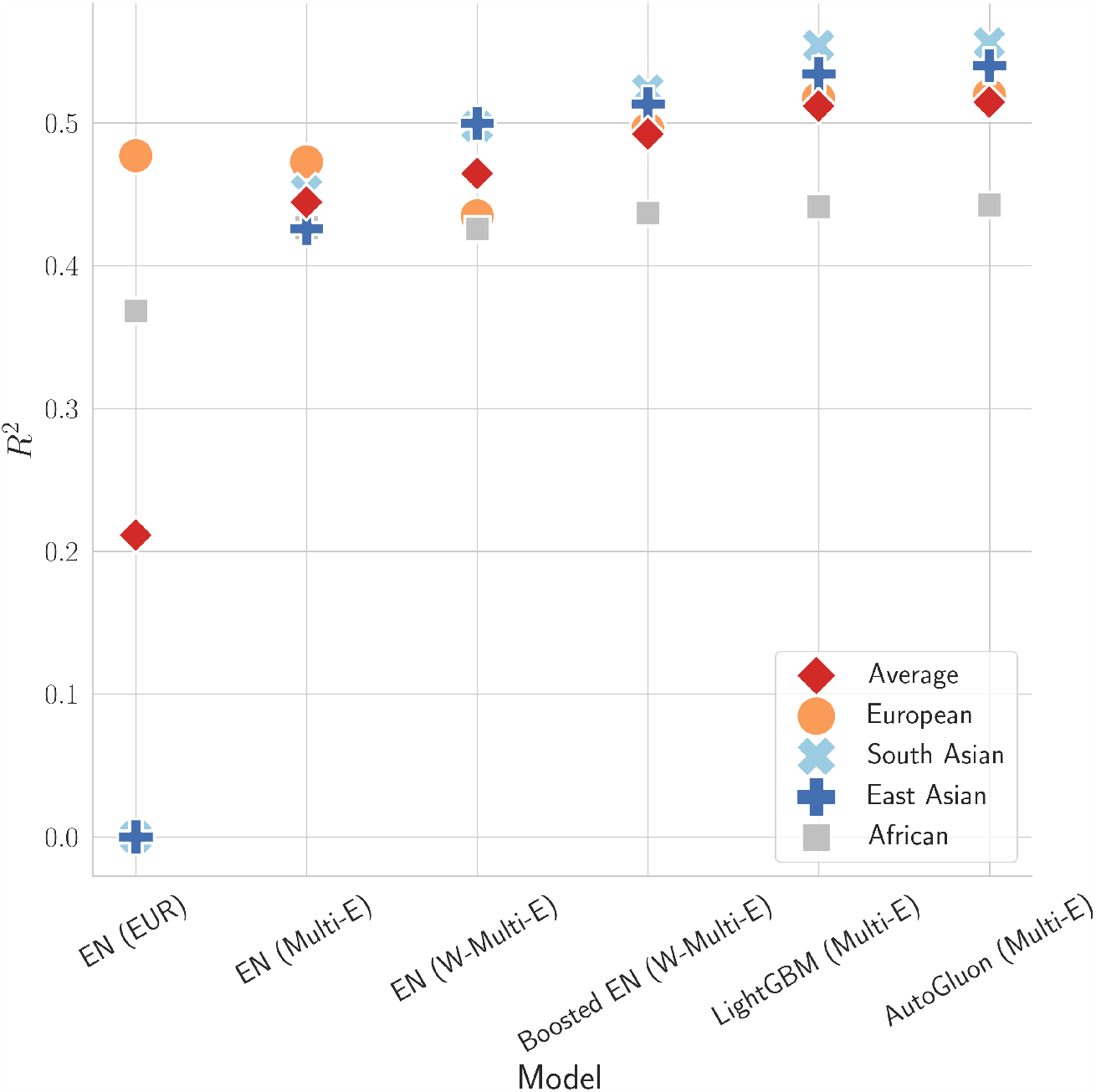
Comparison of *R*^2^ scores across diverse populations for the Standing Height phenotype. “EN” represents Elastic Net. The population used for training is provided in parenthesis, with “EUR” signifying European-only training data, and “Multi-E” indicating the use of multi-ethnic data. The symbol “W” marks the application of population-conditional sample weighting.

We note incremental performance for all populations, starting with Elastic Net which yields an *R*^2^ of 0 for South Asian and East Asian individuals when trained solely on European data. Introducing multi-ethnic data leads to significant *R*^2^ improvements, narrowing the performance gap between populations. Moreover, population-conditional weighting boosts performance for underrepresented groups. Finally, non-linear methods like LightGBM and AutoGluon have proven especially effective for the European, South Asian and East Asian populations. Gains are more modest for the African samples due to the higher genetic variation within this group, making phenotype prediction a more challenging task. Models trained on multi-ethnic datasets can still struggle to capture the intricate relationships between genotype and phenotype specific to African populations. As we integrated increasingly complex and debiasing techniques, we observed an overall improvement in *R*^2^, underscoring that non-linear models, multi-ethnic data, and de-biasing techniques collectively drive enhanced results.

Table 2 provides a comprehensive comparison of various models in predicting standing height across different ancestry groups using diverse training techniques. For the individuals of European descent, training with either European-only or multi-ethnic data showcased similar results, with LightGBM, XGBoost, and AutoGluon emerging as top performers. In contrast, for the South Asian and East Asian groups, introducing multi-ethnic data and applying the proposed population-conditional re-sampling significantly improves predictive performance. The best results in the Asian groups are obtained applying the population-conditional sample weighting with AutoGluon. For the African group, top performance was observed not only with AutoGluon trained on multi-ethnic data but also with the Lasso combined with the population-conditional SMOTE-ENN. This finding underscores the importance of not only model choice but also nuanced training strategies, especially for diverse groups.

**Table 2.** *R*^2^ results for standing height. All the proposed population-conditional (PC) re-sampling methods use multi-ethnic training data. EN: Elastic Net, AG: AutoGluon.

| Population | Training | Lasso | EN | Boosted EN | LightGBM | XGBoost | AG |
| --- | --- | --- | --- | --- | --- | --- | --- |
| European | European-only | 0.508 | 0.477 | 0.506 | **0.520** | **0.520** | **0.520** |
|  | Multi-ethnic | 0.497 | 0.473 | 0.503 | 0.517 | 0.517 | 0.519 |
|  | PC-Random OS | 0.451 | 0.435 | 0.492 | 0.503 | 0.501 | 0.510 |
|  | PC-SMOTE | 0.465 | 0.422 | 0.499 | 0.513 | 0.505 | 0.508 |
|  | PC-SMOTE-ENN | 0.189 | 0 | 0.132 | 0.319 | 0.388 | 0.372 |
|  | PC-Weighted | 0.452 | 0.435 | 0.496 | 0.506 | 0.501 | 0.513 |
| South Asian | European-only | 0 | 0 | 0 | 0 | 0 | 0 |
|  | Multi-ethnic | 0.342 | 0.452 | 0.460 | 0.554 | 0.547 | 0.554 |
|  | PC-Random OS | 0.506 | 0.499 | 0.523 | 0.542 | 0.549 | 0.557 |
|  | PC-SMOTE | 0.486 | 0.480 | 0.509 | 0.552 | 0.533 | 0.541 |
|  | PC-SMOTE-ENN | 0.520 | 0.467 | 0.525 | 0.544 | 0.548 | 0.553 |
|  | PC-Weighted | 0.506 | 0.498 | 0.523 | 0.537 | 0.543 | **0.563** |
| African | European-only | 0.374 | 0.368 | 0.372 | 0.373 | 0.355 | 0.351 |
|  | Multi-ethnic | 0.442 | 0.427 | 0.440 | 0.441 | 0.437 | **0.443** |
|  | PC-Random OS | 0.442 | 0.426 | 0.439 | 0.386 | 0.429 | 0.439 |
|  | PC-SMOTE | 0.434 | 0.411 | 0.421 | 0.400 | 0.433 | 0.414 |
|  | PC-SMOTE-ENN | **0.443** | 0.397 | 0.427 | 0.401 | 0.431 | 0.418 |
|  | PC-Weighted | 0.442 | 0.426 | 0.437 | 0.406 | 0.423 | 0.442 |
| East Asian | European-only | 0 | 0 | 0 | 0 | 0 | 0 |
|  | Multi-ethnic | 0.174 | 0.426 | 0.413 | 0.535 | 0.513 | 0.534 |
|  | PC-Random OS | 0.487 | 0.500 | 0.511 | 0.536 | 0.540 | 0.548 |
|  | PC-SMOTE | 0.466 | 0.479 | 0.497 | 0.525 | 0.526 | 0.547 |
|  | PC-SMOTE-ENN | 0.490 | 0.479 | 0.502 | 0.534 | 0.533 | 0.547 |
|  | PC-Weighted | 0.487 | 0.500 | 0.513 | 0.511 | 0.524 | **0.552** |

### 3.2. Binary phenotypes

We extend our experiments to classification models to observe if they follow similar trends as the regression results presented above. Table 3 showcases the ROC AUC results for two binary phenotypes (diabetes and atrial fibrillation). For both phenotypes, AutoGluon frequently achieves the highest ROC AUC scores, followed by LightGBM, outperforming the linear models. Particularly, the population-conditional weighted training improves model outcomes for the underrepresented groups when using multi-ethnic data.

**Table 3.**
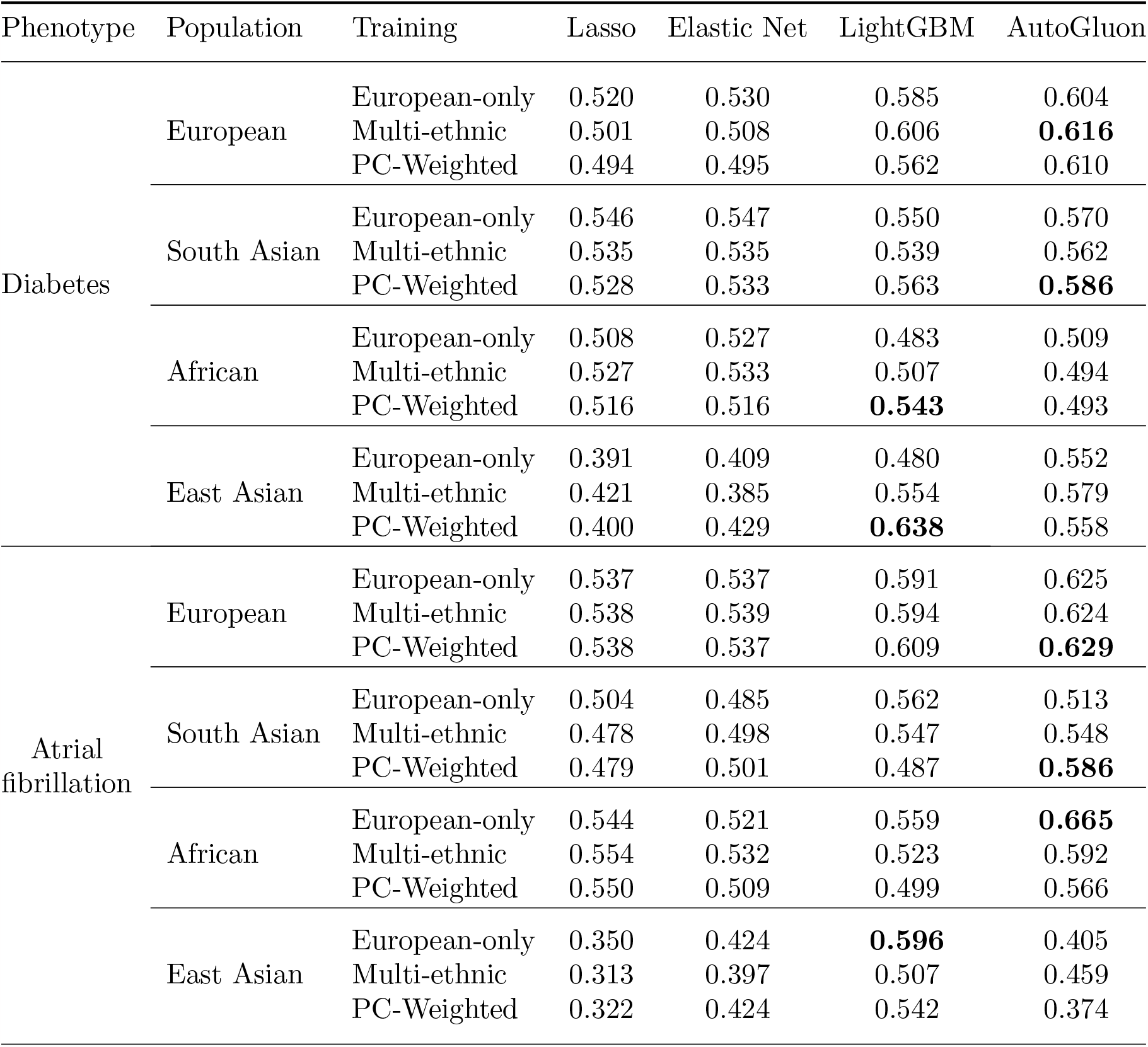
Performance of various models and training techniques in predicting binary phenotypes (Diabetes and Atrial Fibrillation), as measured by ROC AUC scores per group. The proposed population-conditional (PC) method uses multi-ethnic training data.

| Phenotype | Population | Training | Lasso | Elastic Net | LightGBM | AutoGluon |
| --- | --- | --- | --- | --- | --- | --- |
| Diabetes | European | European-only | 0.520 | 0.530 | 0.585 | 0.604 |
|  |  | Multi-ethnic | 0.501 | 0.508 | 0.606 | **0.616** |
|  |  | PC-Weighted | 0.494 | 0.495 | 0.562 | 0.610 |
|  | South Asian | European-only | 0.546 | 0.547 | 0.550 | 0.570 |
|  |  | Multi-ethnic | 0.535 | 0.535 | 0.539 | 0.562 |
|  |  | PC-Weighted | 0.528 | 0.533 | 0.563 | **0.586** |
|  | African | European-only | 0.508 | 0.527 | 0.483 | 0.509 |
|  |  | Multi-ethnic | 0.527 | 0.533 | 0.507 | 0.494 |
|  |  | PC-Weighted | 0.516 | 0.516 | **0.543** | 0.493 |
|  | East Asian | European-only | 0.391 | 0.409 | 0.480 | 0.552 |
|  |  | Multi-ethnic | 0.421 | 0.385 | 0.554 | 0.579 |
|  |  | PC-Weighted | 0.400 | 0.429 | **0.638** | 0.558 |
| Atrial fibrillation | European | European-only | 0.537 | 0.537 | 0.591 | 0.625 |
|  |  | Multi-ethnic | 0.538 | 0.539 | 0.594 | 0.624 |
|  |  | PC-Weighted | 0.538 | 0.537 | 0.609 | **0.629** |
|  | South Asian | European-only | 0.504 | 0.485 | 0.562 | 0.513 |
|  |  | Multi-ethnic | 0.478 | 0.498 | 0.547 | 0.548 |
|  |  | PC-Weighted | 0.479 | 0.501 | 0.487 | **0.586** |
|  | African | European-only | 0.544 | 0.521 | 0.559 | **0.665** |
|  |  | Multi-ethnic | 0.554 | 0.532 | 0.523 | 0.592 |
|  |  | PC-Weighted | 0.550 | 0.509 | 0.499 | 0.566 |
|  | East Asian | European-only | 0.350 | 0.424 | **0.596** | 0.405 |
|  |  | Multi-ethnic | 0.313 | 0.397 | 0.507 | 0.459 |
|  |  | PC-Weighted | 0.322 | 0.424 | 0.542 | 0.374 |

## 4. Conclusions and Future Work

Our results advocate for the implementation of non-linear and ensemble methods, particularly LightGBM and AutoGluon, combined with the proposed population-conditional techniques to enhance genotype-to-phenotype prediction tasks for populations underrepresented in existing datasets. Strategies such as boosting and population-conditional sample weighting and resampling proved to be influential additions in order to better generalize across population and improve prediction accuracy. These methods were effective for both continuous and binary phenotypes, demonstrating their applicability for both regression and classification models.

Our study illustrates the use of methodological advancements to enhance prediction accuracy in the face of a lack of diverse genetic datasets. While the ideal solution would simply be the inclusion of more representative datasets, this is not an accurate reflection of the current data landscape. As such, we recommend for our models and techniques to be implemented when researchers are dealing with datasets of biased representation, especially in genetics. Using these methods should be a priority in situations demanding equitable outcomes, such as in clinical studies.

Failure to address these disparities could engender biases in precision medicine, which might unfavorably impact underrepresented populations. While our study addressed twelve phenotypes, expanding this focus to include other disease phenotypes in future research could yield a deeper understanding of genetic influences on disease. Although AutoGluon includes simple neural network models, future work could delve into a broader spectrum of deep learning architectures, including convolutional layers and attention mechanisms.

The moderate improvement in the African population compared to the Asian groups when applying multi-ethnic training and population-conditional re-sampling can be attributed to the inherent genetic diversity present within the African group, as the SNPs selected for this study are predominantly enriched for representation in Eurasian populations. For future work, a more refined SNP selection tailored for more diverse ancestral backgrounds could potentially enhance the predictive performance and rectify this limitation. A deeper investigation into linkage disequilibrium among SNPs could also optimize the SNP selection process by minimizing redundancies. Although models studied are able to capture genotype-phenotype relationships, covariates, particularly genetic principal components, could allow for a more accurate accounting of the underlying population structure. Incorporating advanced explainable ML techniques^51^ alongside further analysis of covariates can elucidate the underlying mechanisms through which non-linear relationships boost predictive performance, offering a clearer insight into genotype-phenotype mappings. These approaches could refine model performance and enhance prediction accuracy across different ancestry backgrounds.

Given the prevalent bias in many clinical and genetic datasets,^10^ underrepresented populations are often overlooked, with potentially grave implications for health outcomes. This issue is especially pertinent in an era where precision health methods and AI algorithms are becoming increasingly prominent. Thus, implementing strategies such as those presented in our study could considerably enhance the equability and effectiveness of precision medicine for underrepresented groups.

## Acknowledgments

This work was partially supported by NIH under award R01HG010140 and by a grant from the Stanford Institute for Human-Centered Artificial Intelligence (HAI). This research has been conducted using the UK Biobank Resource under Application Number 24983.

Preprint of an article published in Pacific Symposium on Biocomputing © 2023 World Scientific Publishing Co., Singapore, http://psb.stanford.edu/.

